# The voltage-sensing mechanism of the KvAP channel involves breaking of the S4 helix

**DOI:** 10.1101/2019.12.28.889881

**Authors:** Olivier Bignucolo, Simon Bernèche

## Abstract

Voltage-gated ion channels allow ion permeation upon changes of the membrane electrostatic potential (Vm). Each subunit of these tetrameric channels is composed of six transmembrane helices, of which the anti-parallel helix bundle S1-S4 constitutes the voltage-sensor domain (VSD) and S5-S6 forms the pore domain. Here, using molecular dynamics (MD) simulations, we report novel responses of the archaebacterial potassium channel KvAP to cell polarization. We show that the S4 helix, which is straight in the experimental crystal structure solved under depolarized conditions (Vm ∼ 0), breaks into two segments when the cell is polarized (Vm << 0), and reversibly forms a single straight helix following depolarization of the cell (Vm =0). The outermost segment of S4 translates along the normal to the membrane, bringing new perspective to previously paradoxical accessibility experiments that were initially thought to imply the displacement of the whole VSD across the membrane. Our simulations of KvAP reveal that the breaking of S4 under polarization is not a feature unique to hyperpolarization activated channel, as might be suggested by recent cryo-EM structures and MD simulations of the HCN channel.

## Introduction

Voltage-gated potassium channels (Kv) are tetramers that open and close as a function of the membrane electrostatic potential (*1*). Each subunit is composed of six transmembrane helices S1-S6. Voltage dependence is granted by helices S1 to S4, an anti-parallel helical bundle constituting the voltage-sensor domain (VSD), which is linked to the pore domain composed of helices S5 and S6. A much-conserved structural feature of the voltage-sensor domains is a series of four to six basic residues distributed along the S4 helix, each one followed by two hydrophobic residues. The voltage-sensing properties are attributed to these positively charged residues, which are assumed to respond to the membrane electrostatic potential (Vm) by a translation along the membrane normal. This results in an apparent charge transport, or gating current (*2-4*). Under depolarized potential, the pore is open and the channel enters its active state, which can be determined experimentally. The resting or closed state under polarized potentials has been more challenging to investigate.

A “consensus” mechanism describing the voltage-dependent conformational changes of Kv channels in response to variations of the membrane electrostatic potential was developed (*5*), through the integration of several computational studies, notably based on the structures of the Kv1.2 and Kv1.2/2.1 chimera channels (*6-15*). The proposed model consists, within a few Angstroms uncertainty, of a sliding helix mechanism in which S4 undergoes an ensemble of transitions towards a resting state, involving a rotation and a translation along its main axis and toward the intracellular compartment. The translation is ∼ 10Å long, with a spread of 3-4Å. While most of the current knowledge on eukaryotic Kv channels was incorporated in this model, data from the archaea KvAP channel seemed incompatible with the proposed mechanism.

The structure of KvAP voltage-sensing domain was solved by crystallography and NMR (*16-18*), and more recently its complete structure was solved by cryo-EM (*19*). These experiments being performed in absence of any membrane voltage, only the active state of the KvAP VSD could be captured. Its elusive resting state has nevertheless been characterized by several biophysical studies (*20-23*). The sliding helix model depends on the possibility of S4 to exert a translation along its axis in response to variation of the membrane electrostatic potential. As shown in Figure 1A-C, while such a movement is plausible for Kv1.2 and Kv1.2/Kv2.1, there is essentially no room for the long S4 helix of KvAP (33 vs 20 residues) to slide upon depolarization without exposing hydrophobic residues to the polar environment of phospholipid head groups or the solvent.

**Figure 1.**
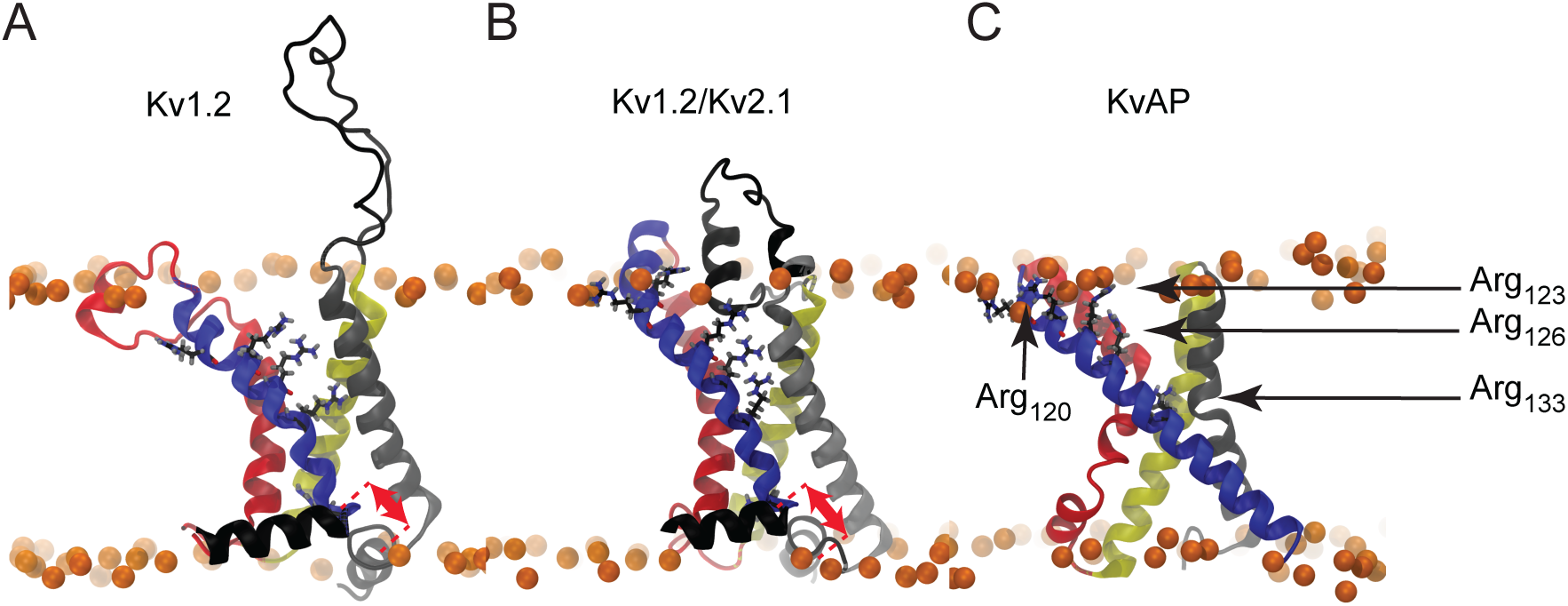
Molecular representations of Kv channel structures. The crystal structures of A) Kv1.2, B) Kv1.2/Kv2.1and C) KvAP are shown embedded in a lipid bilayer (Phosphorus atoms shown as orange spheres). The red arrows illustrate the different translational motions that S4 is likely to undergo without exposing any hydrophobic residue outside of the bilayer core. Such a translation seems not possible in the case of the KvAP crystal structure, thus there is no arrow in panel C. The helices are colored as follows: S1: grey, S2: yellow, S3: red, S4: blue. The arginine residues of the S4 N-ter are shown as sticks with carbon, nitrogen, oxygen in black, blue and red.

One key experimental observation, for which a mechanistic explanation remains elusive since its publication, is the accessibility measurements of avidin to biotinylated cysteins in the S3 and S4 segments (*21, 23*). These experiments showed that avidin in the intracellular space could bind to biotinylated cysteine located in the middle of S4. This observation was explained by a large displacement of the whole S4 helix across the hydrophobic core of the membrane, which was difficult to reconcile with other experiments (*24*).

Our unrestrained simulations show that, under cell polarization, the voltage-sensing domain of KvAP undergoes a transition that involves the rupture of the Asp_62_-Arg_133_ salt bridge immediately followed by the formation of a kink in the middle of S4. The resulting sliding movement of the kinked S4 helix towards the intracellular space provides a more consensual explanation to the avidin accessibility experiments. Similar conformational changes were recently observed in the VSD of HCN, a channel activated by hyperpolarization (*25, 26*). The breaking of S4 observed in the VSD of KvAP reveals that this transition is not specific to hyperpolarization gating. Sequence alignment reveals that the specific sequence of the KvAP S4 N-ter is hardly found in any eukaryotic voltage-gated ion channel (see Table 1). However, this sequence is found in several archaea and prokaryotes, among which many pathogens, making it a potential selective target for antibiotic investigations.

**Table 1.**
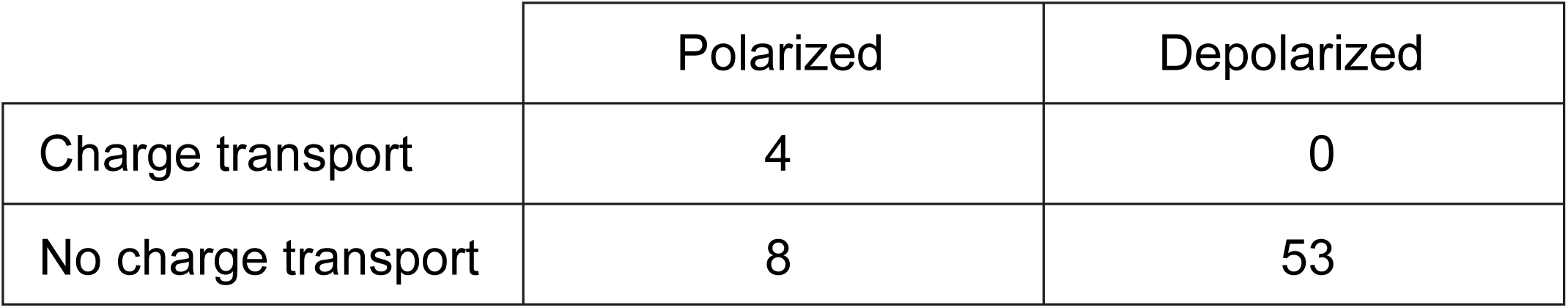
Summary of the charge transport events observed during this study.

|  | Polarized | Depolarized |
| --- | --- | --- |
| Charge transport | 4 | 0 |
| No charge transport | 8 | 53 |

## Results

### KvAP response to cell polarization involves bending of S4

To address the ill-defined mechanism of voltage-sensing in prokaryotic cells, we carried out a large number of independent molecular dynamics (MD) simulations in which the VSDs were exposed to a wide range of membrane electrostatic potentials. As shown in Figure 2, a system consisted of two bilayers mimics a cell with two separated water compartments that, according to the orientation of the bilayer leaflets and incorporated proteins, correspond to the extra- and intracellular compartments. This compartmentalization allows one to adjust the membrane potential by changing the number of ions in either compartment (*27-29*). We constructed 66 such systems, allowing the study of 132 VSDs. The simulated membrane potential (Vm) ranged from -1.7 to 0.5 V. While these Vm values are of higher magnitude than physiologically found in cells, they remain in a range that does not expose the membrane to electroporation.

**Figure 2.**
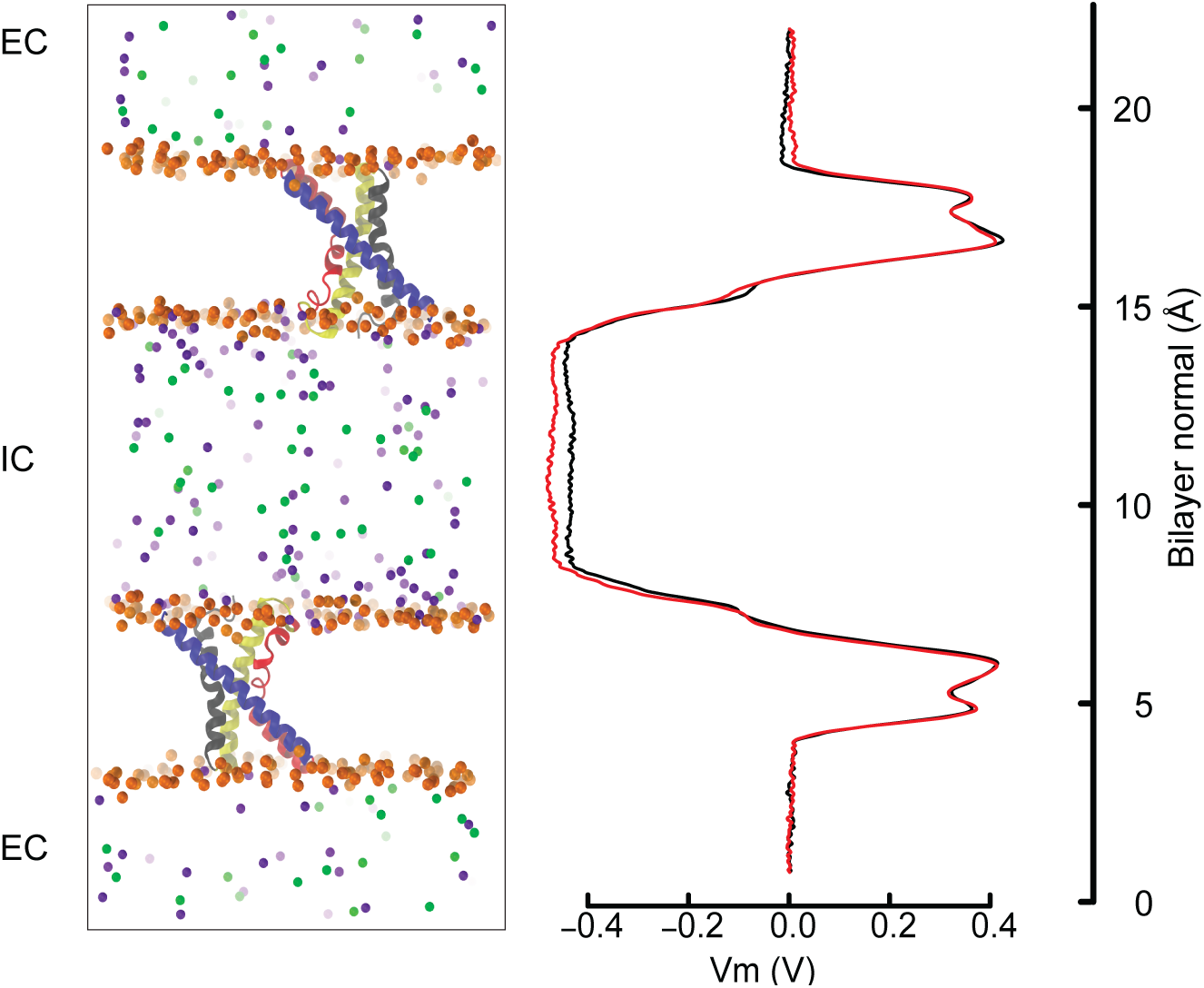
The double bilayer system enables the explicit tuning of the membrane potential. Left) Molecular representation of two antiparallel bilayers with embedded VSDs. The proteins are represented and colored as in Figure 1 and the phosphorous atoms as well as the ions are depicted as spheres. Phosphorous, sodium and chloride are colored orange, violet and green. For clarity, water and lipid molecules have been removed. All the atoms shown constitute one simulated entity. Because of the periodic boundary conditions, the water and ions above and below each displayed bilayer communicate and form a single slab (EC: extracellular), which is isolated from the middle slab (IC: intracellular). Right) The electrostatic potential along the bilayer normal (z) is shown adjacent to the molecular representation. We draw the potential at the beginning (0-20 ns, black) and at the end (last 20 ns, red), which is normalized to 0 V at z = 0, i.e. in the E.C. The similarity of the two curves indicates that the charge imbalance manually set between the two compartments remained stable during the simulation.

We controlled the electrostatic steadiness of the bilayer systems by monitoring the difference ***Δ***Vm between the membrane potential averaged over the first and last 20 ns of the trajectory (see Figure 2). As a consequence of the limited size of the systems, which typically contain ∼ 240 000 atoms, a single charge transport across the membrane induces a ***Δ***Vm of ∼ 200 mV. In four of the 66 simulations we detected variation of membrane potential that indicated charge transport equivalent to the relocation of one or two ions across the membrane. As shown in Table 1, these events occurred only when the membrane was polarized (negative potential) (Fisher’s exact test probability p = 0.007).

The crystal structure of KvAP is characterized by two funnels readily accessible to the solvent, as can be seen in Figure 1C. An important salt bridge (figure 3) between Asp_62_ (helix S1) and Arg_133_ (helix S4) in the middle of the membrane constitutes the only barrier between the extra- and intracellular compartments. We observed the rupture of this salt bridge in all simulations in which a charge transport occurred (Table 1, Figures 3 and 4), and in none of the others. Upon rupture of the salt bridge, the negative charge of Asp_62_ moved toward the extracellular compartment, while the positive charge of Arg_133_ moved toward the intracellular compartment, which resulted in the observed gating charge transport. In addition, whereas the S4 helix is straight under depolarized conditions, it formed the evoked kink at the level of Gly_134_ only when the cell was polarized and the salt bridge was broken. Consequently, S4 was split in two segments, the one on the intracellular side being reoriented in a direction almost parallel to the membrane surface (Figure 3, right), like the S4-S5 linker of Kv1.2 and Kv1.2/Kv2.1 (Figures 1A and 1B).

**Figure 3.**
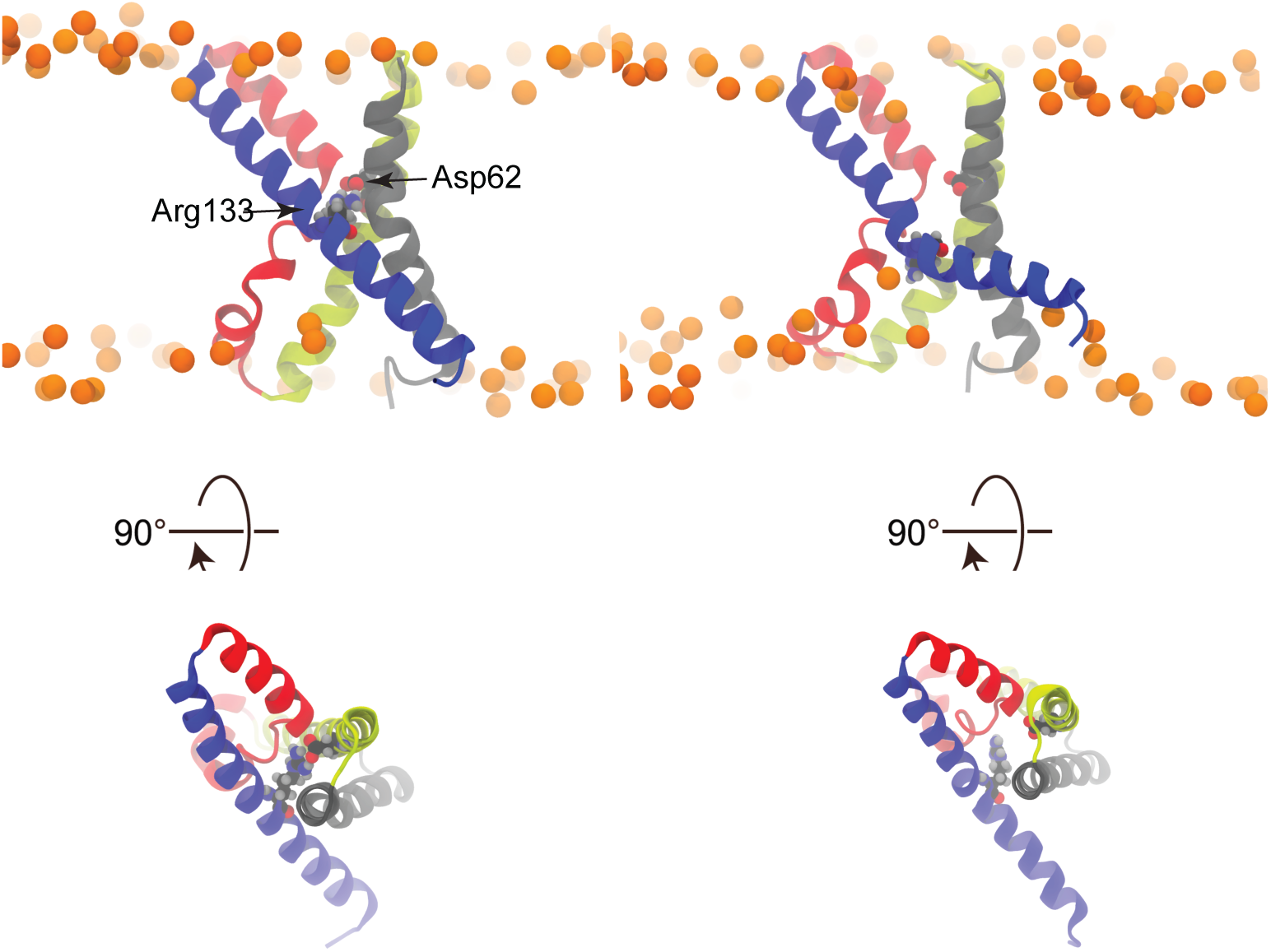
Salt bridge rupture upon polarization. A) S4 is straight and the Asp_62_-Arg_133_ salt bridge is intact in the X-ray structure. B) Representative snapshot from a simulation conducted under polarized potential, taken after the charge transport, showing kinked S4 and broken salt bridge.

**Figure 4.**
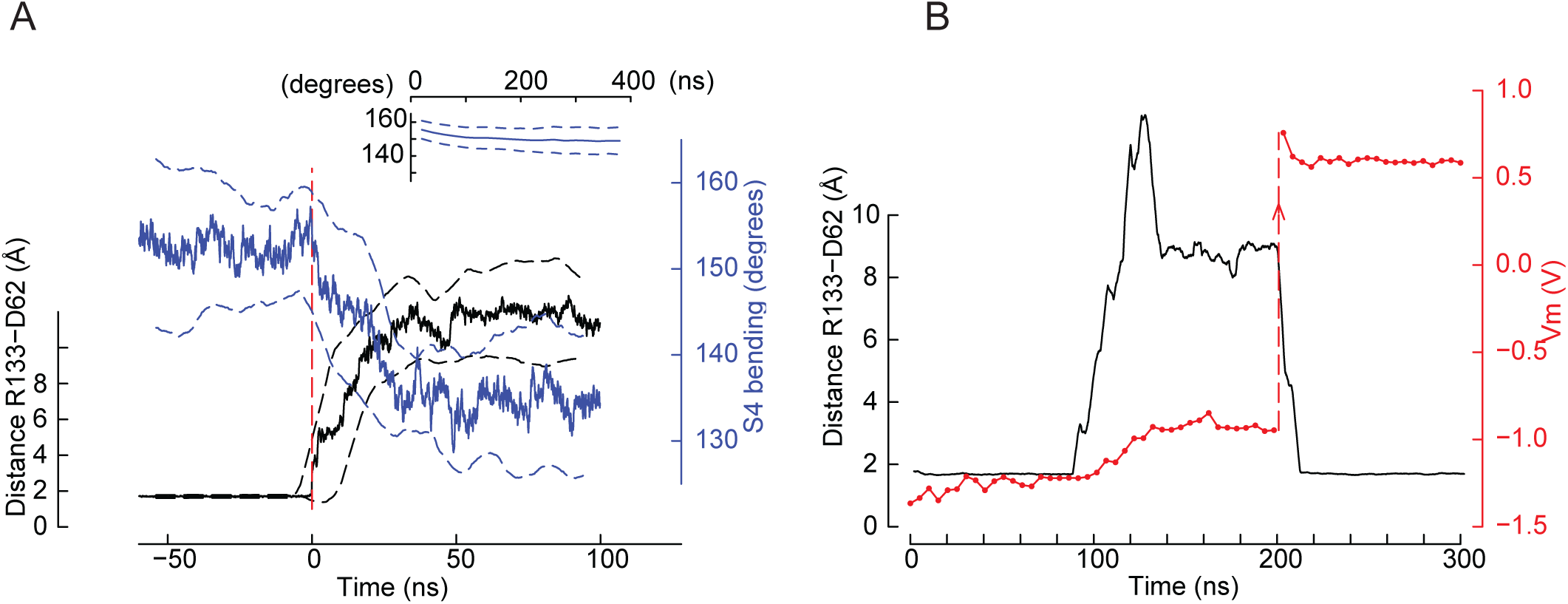
Co-occurrence of the salt bridge rupture and the kink formation in S4 upon polarization. A) Time series of the minimal distance between Asp_62_ and Arg_133_ (black, left y-axis) and the bending of S4 (blue, right y-axis). Mean ± standard deviations from four pooled independent trajectories are shown. Before pooling, the time axis of each trajectory was set to 0 when the Asp_62_ - Arg_133_ distance goes above 3 Å for the first time (red dashed vertical line). For comparison, the inset shows the S4 angle time evolution (Mean ± standard deviation) of trajectories conducted at Vm = 0 V (n = 62). B) The status of the Asp_62_-Arg_133_ salt bridge depends upon the membrane potential. Black lines, left y-axis: distance between Asp_62_ and Arg_133_ as a function of simulation time. Red lines, right y-axis: membrane potential as a function of simulation time. At t∼90ns, rupture of the salt bridge. At t=200, Vm manual sign inversion by ion displacement (red arrow). At t∼210ns, restoration of the salt bridge.

A time series analysis of the trajectories in which the gating charge transport occurred shows that the breaking of the salt bridge and the kink in S4 occur concurrently (Figure 4A). None of these events was observed in the simulations conducted at Vm ∼ 0 V (Figure 4A, inset). This time series analysis supports the ideas that 1) the two conformational changes are related and 2) they are due to the membrane polarization. We further asked whether the membrane potential alone drives the status of the Asp_62_-Arg_133_ salt bridge. In a particular simulation initialized under a strong polarizing potential, the salt bridge broke after ∼ 90 ns. Consecutively to the reorientation of the Asp_62_ and Arg_133_ side chains, the membrane potential decreased within ∼ 30 ns to a value corresponding to a gating charge transport of two units and remained stable during the next 50-60 ns. With the aim of testing the dependence of the conformational changes on the membrane potential, we stopped the simulation and moved ions between the extra- and intracellular compartment in order to mimic a depolarized potential. We then continued the simulation from this new starting point. Within ∼10 ns of simulation, the side chains of Asp_62_ and Arg_133_ reoriented and restored the salt bridge, which remained intact for the next 200 ns (Figure 4B).

Interestingly, experimental (*17, 18, 20, 30*) and computational (*31*) investigations have reported either that the S4 of KvAP may kink near the middle of the bilayer, or that the Asp_62_- Arg_133_ salt bridge may break (*32*). In an NMR structure determination of the KvAP VSD, a loss of helical periodicity was identified at the level of Gly_134_, suggesting that the S4 helix might be constituted of two helices connected by a hinge comprising Ile_131_, Ser_132_ and Arg_133_ (*18*). Interestingly, three of the 20 conformations deposited for the KvAP VSD NMR solution structure (code 2KYH) display a kink in the middle of S4 (*17*).

Whereas experimental or computational studies support the idea of a hinge in the middle of S4 or the rupture of the salt bridge, the present study shows for the first time that these two conformational changes happen simultaneously upon polarization and that they lead to the charge transport observed experimentally and generally interpreted as a gating current.

### Avidin binding to biotinylated KvAP voltage-sensor domain

In 2003 and 2005, Jiang et al.(*23*) and Ruta et al.(*21*) reported experiments in which the binding of avidin to biotinylated cysteines was used to deduce the residue accessibility from the external or internal cell compartments. In the study described by Jiang et al., a 17Å linker connected the Cys Cα atom to biotin through an amide bond. These avidin binding experiments notably showed that the biotinylated residues 125 and 127, located in the upper half of the S4 helix, were accessible to avidin from the intracellular side. These results supported the idea of a voltage-sensor paddle model in which the helix-turn-helix S3a-S4 moves through the membrane upon voltage changes. However, this model requires an important movement of charged amino acids across the hydrophobic core of the membrane, which was difficult to reconcile with other observations(*24*).

Our MD simulations show that the bending of S4 induced by membrane polarization reduces sufficiently the distance required for avidin to bind to the biotinylated residues without requiring the S3a and S4 helices to cross the membrane core. In order to determine how close an avidin molecule could come to the residues investigated by Jiang et al. (*23*) and Ruta et al. (*21*) considering the conformational change of S4 under membrane polarization, we conducted a steered molecular dynamics (SMD) investigation involving the KvAP VSD and the monomeric avidin-biotin complex (Figure 5A). A soft constant pulling force (see Methods) was exerted between the biotin carboxylic acid functional group and the Cα of Ile_127_. We performed the steered molecular dynamics simulations (SMD) starting either from the X-ray structure or from a conformation displaying the kink in S4 described above, with an initial distance between the Cα of Ile_127_ and the biotin carboxylic acid of ∼25-26 Å. Whereas this distance stabilized to ∼19 Å during 50 ns of constant pulling in the case of a VSD with a straight S4 helix, it decreased steadily, attaining ∼ 11Å, and stabilized to a value of ∼13 Å in the case of the kinked S4 helix (Figure 5). This distance is significantly less than the experimentally used linker, and thus the response of the VSD to membrane polarization is compatible with the accessibility experiments described above.

**Figure 5.**
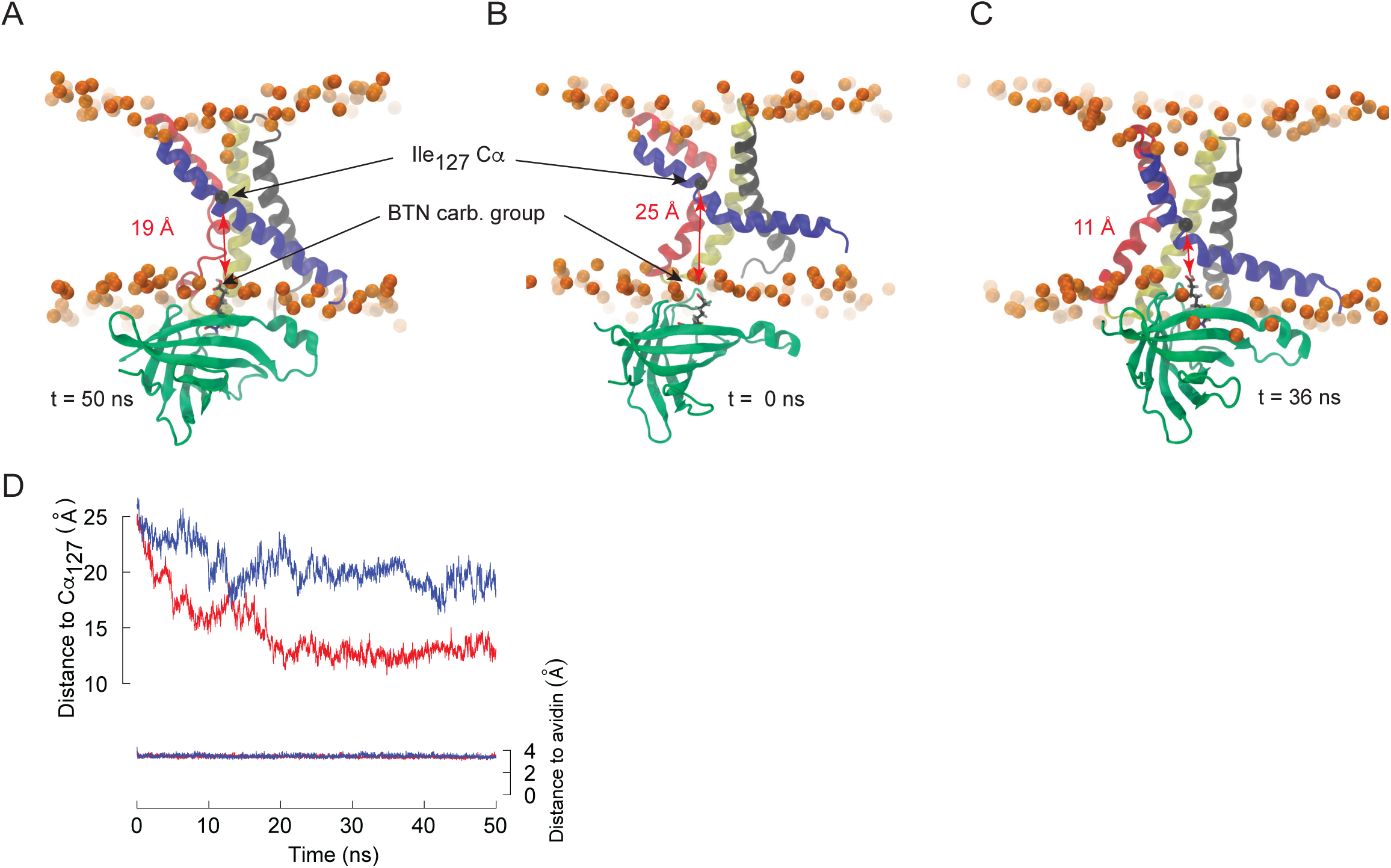
The kink in S4 allows binding of avidin to biotinylated residues in the middle of S4. Snapshots extracted from steered molecular dynamics simulations initiated with a VSD in which the S4 helix is straight (A: t = 50 ns) or kinked (B: t = 0 ns, C: t=36 ns). Avidin is colored in green, and biotin is shown in stick representation with carbon black, oxygen red and hydrogen grey. The distance between biotin and the Cα (black sphere) of ILE_127_ is highlighted. D) Time series of the distance between the carboxylic group of biotin and the Cα of ILE_127_ for the simulations with KvAP as in the crystal structure (blue) or harboring the kink in S4 (red). For control, the distance between the biotin and its binding site to avidin is shown (right y axis). Pulling forces were applied between the biotin and the ILE_127_ Cα, and between the biotin and the avidin binding site (see Methods).

## Discussion

This study reveals a novel response of KvAP to cell polarization that consists in the concurrent disruption of the salt bridge between Asp_62_ in S1 and Arg_133_ in S4 and the formation of a kink at the level of Gly_134_ in S4. Whereas the rupture of the Asp_62_-Arg_133_ salt bridge has been reported as a response to polarization (*33*), the current study shows that the rupture of the salt bridge induces the formation of a kink in S4. The induced conformation is expected to further allow the translation of the upper part of S4 along its principal axis and towards the intracellular side of the bilayer.

The helix breaking occurs at the level of a Gly residue located 8 positions downstream from the S4 basic residues. Sequence alignments reveal that the specific features of the KvAP S4 sequence are not seen in any eukaryotic voltage-gated ion channel, whereas there are found in several archaea and prokaryotes, among which many pathogens, as further explained in Supplementary Material.

The observed conformational changes imply that a tethered biotin on the external half of S4 is accessible to avidin from the intracellular compartment, bringing a biophysically coherent explanation to the accessibility experiments described by Jiang et al (*23*), and Ruta et al (*21*). Studies involving both the voltage-sensor and the pore would be required to further investigate the postulated function of the lower half of S4 as a surrogate S4-S5 linker, notably in the non-domain-swapped architecture revealed by a recent cryo-EM structure of KvAP (*19*).

In a recent cryo-EM study on the HCN channel, a hyperpolarization activated potassium channel, a disulfide bridge was generated between F186C in helix S2 and S264C in helix S4 with the aim of mimicking hyperpolarized conditions. Consequently, the VSD was trapped in a presumably activated state, characterized by a kink in S4 at the level of the disulfide bridge and a sliding movement of the external part of S4 towards the intracellular side (*25*). Similar conformational changes were observed in MD simulations of the HCN voltage-sensor domain under membrane hyperpolarization (*26*). The breaking of the S4 helix in two smaller helices was suggested to be essential to hyperpolarization gating. Our unrestrained simulations reveal that this feature is not unique to channel activated by cell hyper polarization, but is also observed in a channel activated by depolarization, like KvAP.

## Methods

### Sequence analysis

The non-redundant UniProt/SwissProt sequence database was used for searching voltage-gated potassium channels homologs to the KvAP VSD sequence, ID Q9YDF8 (*34, 35*). The sequences were further curated using in-house Python scripts in order to remove undefined species, uncharacterized fragments, retain sequences of length similar to the KvAP VSD ± 100 residues. The scripts further selected sequences characterized by the typical feature of a voltage-sensor domain, i.e. a series of three triplets consisting of a pair of mostly hydrophobic residues followed by a basic residue and, in addition, a segment of seven any residues followed by a Gly. This last criterion allowed us to discriminate the sequences according to the specificity of the S4 helix described in this work.

### MD simulations

The atomic model of the KvAP VSD was based on the crystal structure PDB code 1ORS, assumed to correspond to the active state of the channel (*36*). The structure of a complete KvAP channel was solved recently through cryo-EM (*19*). Note, however, that the structure of the voltage-sensor domain is identical in the previous and new structures (see suppl. Figure 2). Specifically, S4 is straight in both structures, which were solved at Vm = 0V. The VSD was inserted in an asymmetric bilayer using the CHARMM-GUI web service (*37*). The “extra-cellular” leaflet was composed of 100 POPC (1-palmitoyl-2-oleoyl-sn-glycero-3-phosphocholine) and 80 cholesterol, and the “intra-cellular” leaflet was composed of 50 POPC, 50 POPS (1-palmitoyl-2-oleoyl-sn-glycero-3-phosphoserine) and 80 cholesterol molecules. The system was further solvated with ∼ 25,000 water molecules, represented by the TIP3P model (*38*). Neutralizing K^+^ and Cl^-^ counterions were added to mimic a salt concentration of 0.15M. Two such systems were combined in an antiparallel way to form a double bilayer system, simulating a cell membrane separating two different water compartments (*28, 39, 40*). The system contained ∼ 235,000 atoms. The construct contains in its center a water slab simulating the “intracellular” compartment, and the two slabs on the edges are combined through periodic boundary conditions to form the “extracellular” compartment. Using this construct, the membrane potential can be adjusted to the desired value by changing the number of ions in either compartment.

The simulated membrane potential (Vm) ranged from -1.7 to 0.5 V. These high values allowed to augment the conformational space exploration, however without attaining potentials that would induce electroporation. Thus, in a study investigating the stabilization effect of cholesterol on lipid membranes, Casciola *et al.* (*41*) exposed bilayers composed of 0 to 50 mol% cholesterol to membrane potentials values up to 5.35 V. They observed within the first 60-70 ns of simulation that the “electroporation thresholds increased from ∼ 2.3 V for bare bilayers to ∼ 4.4 V as the cholesterol content reached 30 mol% concentration”. In another study, an electroporation thresholds of -1.8 V was reported for a cholesterol free membrane(*42*). According to these data, the membrane potential applied in our work is not expected to destabilize the bilayers, which contain ∼ 45 mol% cholesterol. We effectively did not observe any strong membrane deformation during the simulations, generally of ∼ 200 ns length, reaching 740 ns in one case.

All-atom MD simulations were performed with the GROMACS software package version 4.5(*43*), with the CHARMM force-field (*44*), versions v27 for proteins(*45*) and v36 for lipids(*46*). A constant pressure of 1 bar was maintained using the Berendsen algorithm (time constant 1ps) (*47*). The temperature was kept at 310 K by a stochastic rescaling of the velocities (time constant 0.2 ps) (*48*). Bond lengths and angles involving hydrogen atoms were constrained using the LINCS algorithm(*49*), allowing an integration time step of 2 fs. Short-range electrostatics were cut off at 1.2 nm, and the particle mesh Ewald method was used for long-range electrostatic (*50*). Van der Waals interactions were described with Lennard-Jones potential up to a distance of 1.2 nm. The systems were equilibrated following the CHARMM-GUI protocol(*51*). Independent simulations were conducted on 66 double bilayer systems, thus allowing the study of 132 voltage-sensor domains at various membrane potentials.

For the study of the avidin accessibility, we reasoned that whereas avidin generally forms a tetramer in solution, with extremely high affinity to biotin (K_d_ ∼ 10^−15^ M), it was shown that the monomeric avidin also binds biotin with high affinity (K_d_ ∼ 10^−7^M), which is sufficient to explain the experiments mentioned in the main text(*52*). The access of a monomeric avidin to a VSD-bound biotin is structurally less constrained than that of a tetramer. Consequently, chain A from the complex avidin-biotin crystal structure (PDB code 1AVD) was placed manually at a few angstroms from a bilayer containing KvAP, which was in either the crystallized conformation or after formation of the kink in S4 (Figure 5B shows the initial positions of the molecules in this case). Force field parameters for biotin were generated using the CGenFF web-service (*53*) based on a structure obtained from the Zinc12 database (*54*). Constant pulling forces of 100 kJ•mol^-1^•nm^-1^ were applied between the Cα atom of Ile_127_ of KvAP and the center of mass of the biotin carboxylic acid group. An equal force was applied between the center of mass of the nitrogen and sulphur atoms of biotin and that of Trp_70_ and Trp_97_ of avidin, as these two residues define the biotin-avidin binding site(*55*).

## Author Contributions

O.B. and S.B conceived the study. O.B. performed the numerical simulations and the analytic calculations. O.B and S.B wrote the manuscript.

## Acknowledgments

This work was supported by a grant from the Swiss National Science Foundation (SNF Professorship No PP00P3_139205 to S.B.). Simulations were performed at the Swiss National Supercomputing Centre (CSCS) under projects ID S545 and SM09, and data analysis at sciCORE, the scientific computing center from the University of Basel (http://scicore.unibas.ch/). The authors thank Annaïse Jauch and Niklaus Johner for helpful comments on the manuscript.

## Supplementary Material

### Sequence analysis suggests a potential target for antibiotics research

The specific features of the KvAP S4 N-ter sequence is not seen in any eukaryotic voltage-gated ion channel. The Gly residue at which the observed kink occurs is located 8 positions downstream from the S4 basic residues. As shown in Supplementary Table 1, the most resembling eukaryotic sequences contained a much shorter segment, generally 3 residues, connecting the last S4 basic residue and the next Gly. However, we found that several prokaryotic potassium voltage-gated channels displayed a similar sequence as the KvAP S4 N-ter. Many of them belong to the anaerobic Bacteroides species, which are of significant clinical relevance. *Bacteroides fragilis* infections display a mortality of 20%, which rises to more than 60% if left untreated (1). A study on bacteremia, reveals that while the incidence of anaerobic bacteremia is relatively low (less than 3%), the associated mortality is higher than 20% (2). In the same line, *Bacteroides pyogenes* causes serious human wound infections(3, 4) whereas *Bacteroides thetaitoamicron*, which can exacerbate *Escherichia coli* (E.coli) and *Clostridium difficile* infections, is the second most common infectious anaerobic gram-negative bacteria(5). Thus, the prokaryotic specific mechanism identified in the current study may be used as a selective target addressing several lethal pathogens, some of them being a cause for major concern in the context of increasing reported cases of antimicrobial (β-lactam, Carbapenems and other antibiotics) resistance (6-9). Mutation experiments involving the *E. Coli* Kch potassium channel suggested that it maintains the membrane potential and could prove essential under certain stress conditions, like higher external potassium concentration(10). Further, in support to the pertinence of potassium channels as potential antibacterial targets, gastric colonization by *Helicobacter pylori* lacking its potassium channel HpKchA is impaired (11).

### Ion and water transport

In two instances, we observed a potassium ion moving from the extracellular to the intracellular compartment, as well as several water molecules. In a previous molecular dynamics study, Freites et al. reported the formation of a water pore through the KvAP VSD, pointing the Asp_62_- Arg_133_ salt bridge rupture as a requirement for water transport (14). However, we propose that this opening is rather transient, and that a further sliding of S4, though not observed in our rather short simulations, would allow Arg_133_ to form new interactions with acidic residues located more toward the intracellular side, namely Asp_72_ in S2 and Glu_93_ in S3. These new interactions would then restrict the transient opening through the VSD.

### The pore domain and the membrane

In this study, only the voltage sensor of KvAP is included in the simulation systems, without the pore domain. The conformation and dynamics of the isolated VSD is expected to be different from that observed in presence of the pore domain. Nevertheless, the polarization is the driving force inducing the disruption of the salt bridge and the displacement of the positively charged upper segment of S4. The direction and magnitude of this force does not depend on the fact that the VSD is linked or not to the pore domain, and thus these two events are expected to happen in the case of a whole channel as well. In our simulations, the intracellular segment of S4 moved unconstrained, with a tendency to orient parallel to the membrane. In principle, this movement could be different in the case of the whole channel. The comparison of the whole chain structure and the structure of the voltage sensor alone shows, however, that the conformation of the S1-S4 helices in the open state is not affected by the pore domain (Suppl. Figure 2).

We noticed a narrowing of the lipid bilayer in the vicinity of the protein. Even residues Arg_123_ and Arg_126_, which are located relatively deep towards the middle of the bilayer, formed hydrogen bonds with the lipids. The water filled cavities, on both sides of the membrane, also induced rearrangements of the phospholipids. As shown in Figure 3B, a few phosphate groups entered in the water filled cavity, although they generally remained at the most external part of it. We wondered whether the formation of the kink, which may further enlarge the water filled cavity on the intracellular side, would affect the structure of the bilayer. As shown in Supplementary Figure 1, the intracellular leaflet thickness was ∼ 20 Å resp. 17 Å for lipids situated far from resp. near the VSD. However, the thinning of the bilayer was the same whether S4 formed a kink or not. To be more precise, we further plotted the thickness of the membrane as a function of the bending of S4. We also analyzed the thickness of the lower leaflet as a function of the height of the last residue of S4, defined as the distance along the normal to the bilayer between the C*α* atom of Leu_148_ and the average height of the C316 atoms of the lipids, representing the center of the bilayer. In both analyzes (data not shown), there was no relationship between the thickness of the leaflet and the shape of S4, which demonstrates that the kink did not affect the penetration of the lipid head groups into the membrane. On the other hand, the polar head groups tended to reorient in the vicinity of a transmembrane protein. We then wondered whether the formation of the kink in S4 would modify the polar head orientation. The orientation of the phosphocholine head groups is defined by the angle between the Phosphorus to Nitrogen vector and the normal to the bilayer. The average value of this angle was ∼ 71 degrees resp. ∼ 64 for lipids situated far from resp. near the VSD. The kink in S4 has no impact on the reorientation of the lipid polar heads. It is thus concluded that whereas the membrane structure was modified in the vicinity of the channel, these rearrangements were not affected by the disruption of the Asp_62_-Arg_133_ salt bridge and the kink of S4.

---

**Supplementary figure 1.**
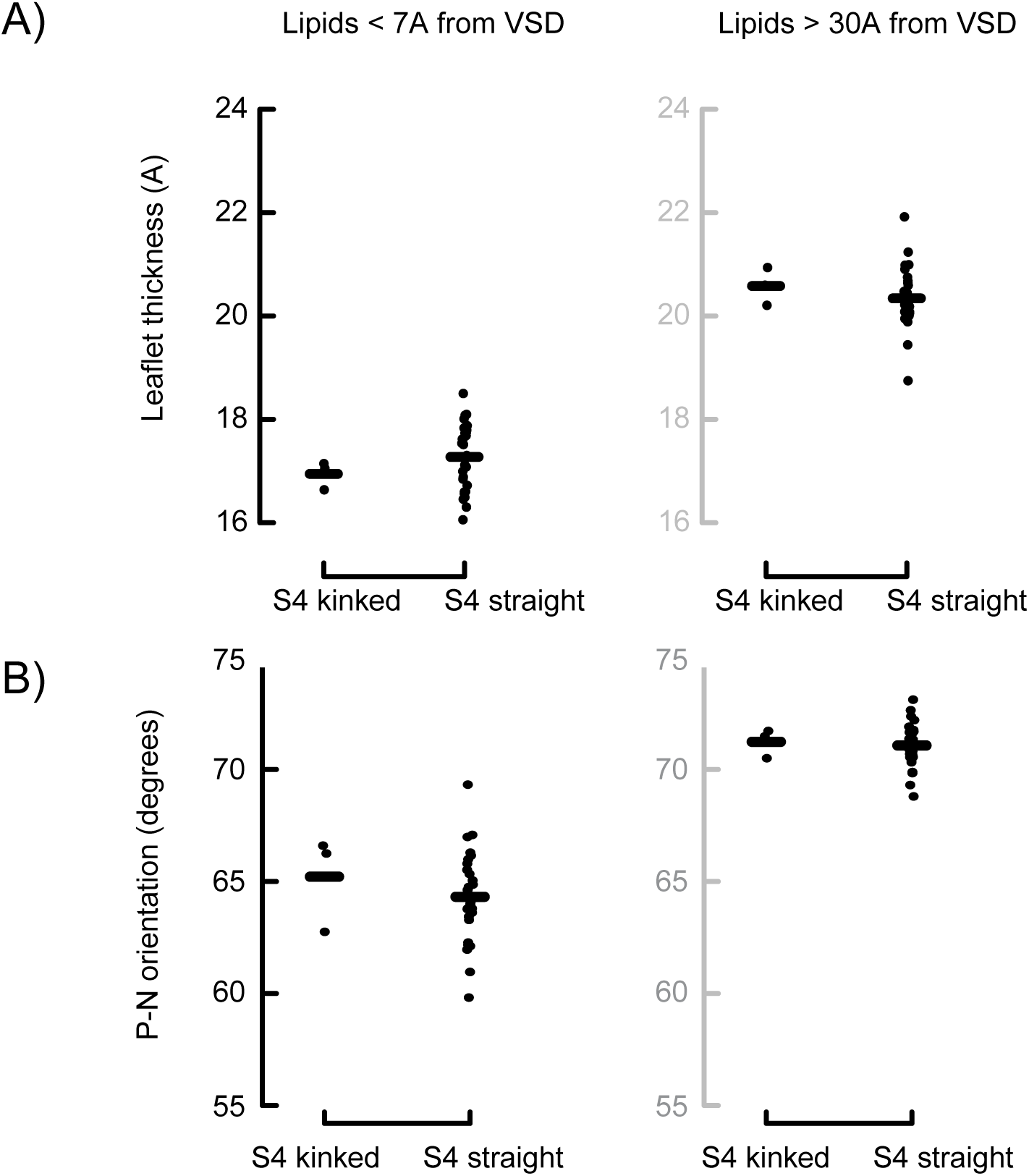

**Supplementary figure 2.**
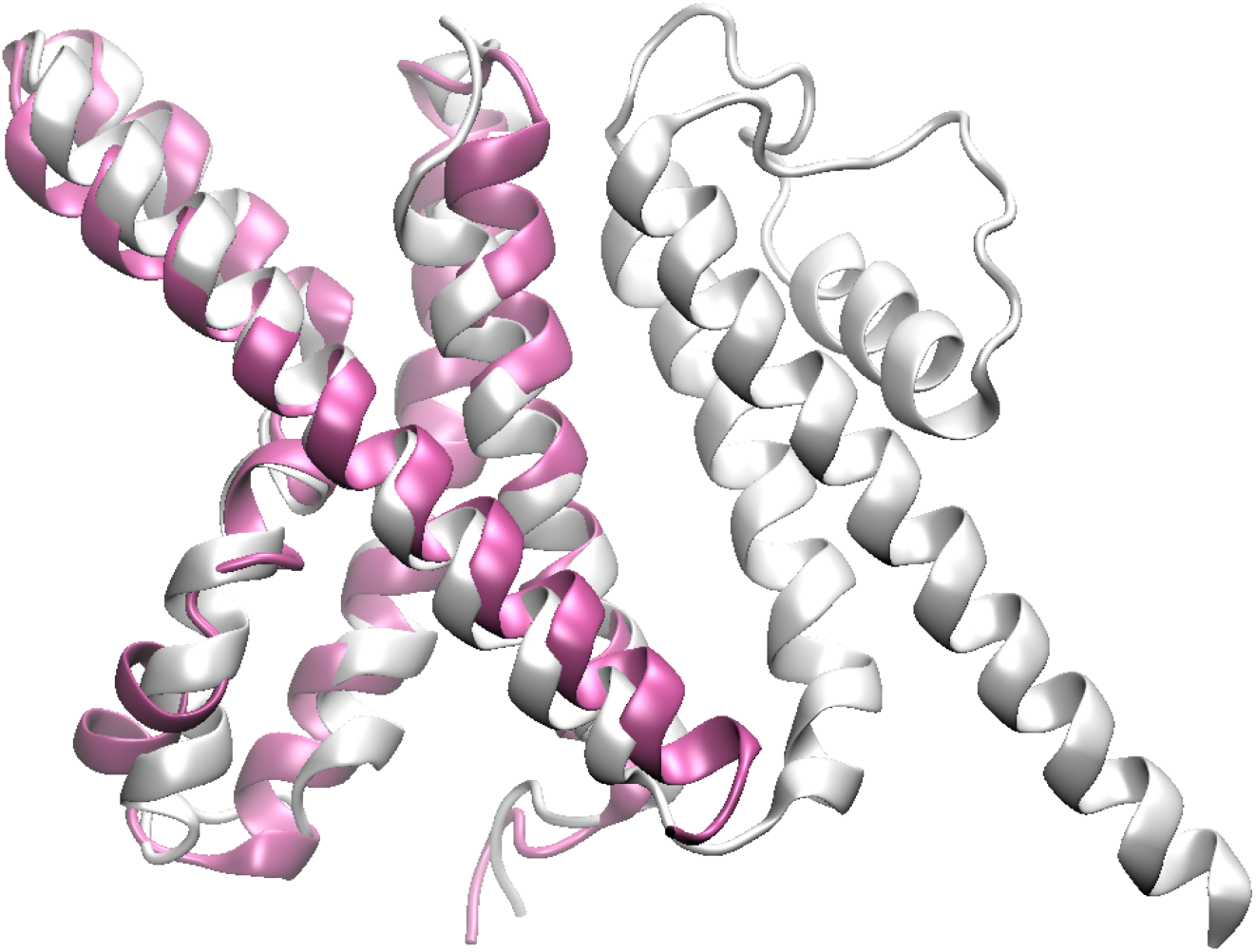

**Supplementary Table 1.**
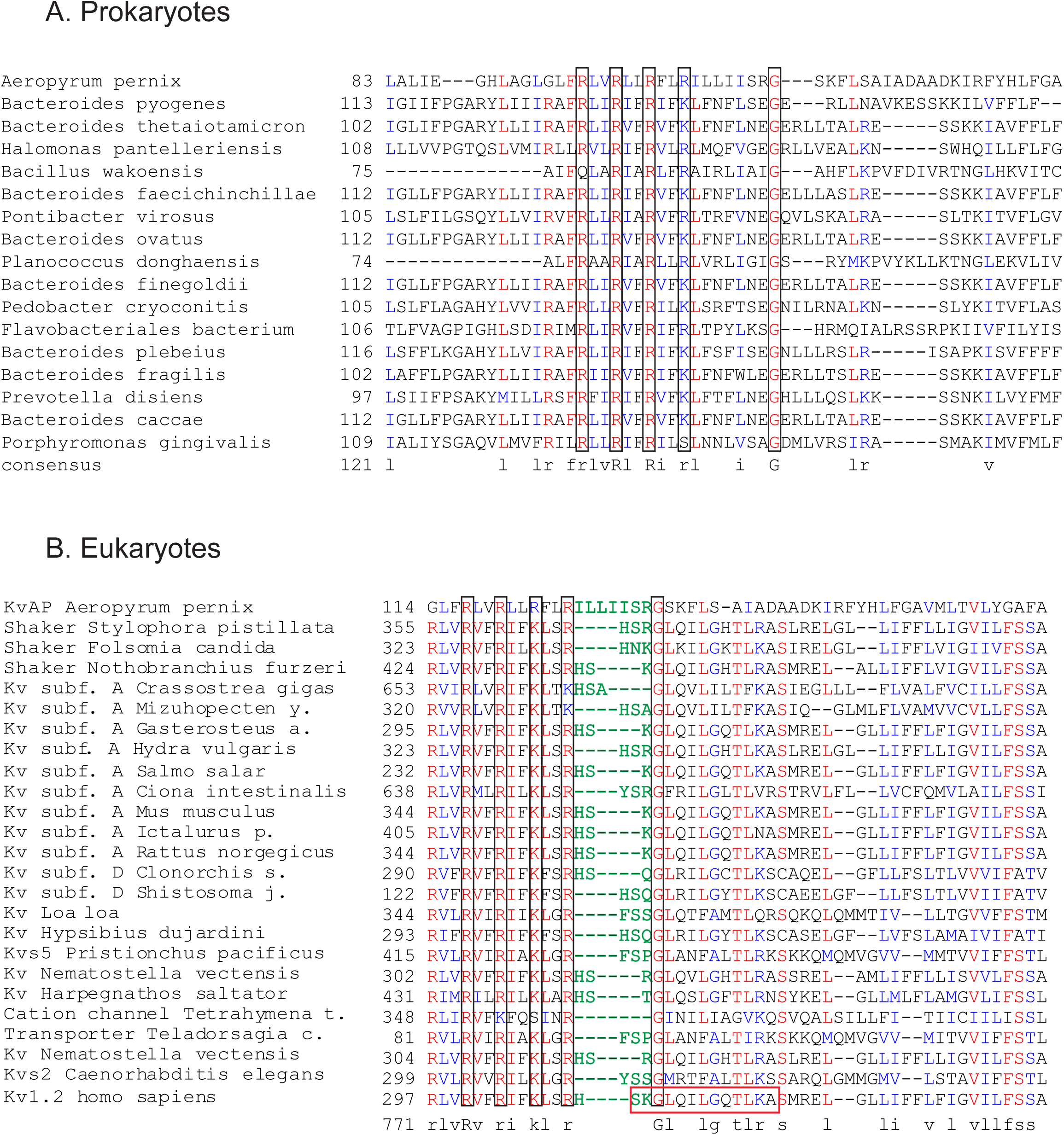

## References

1. K. J. Swartz, Sensing voltage across lipid membranes. Nature 456, 891–897 (2008).

2. Q. Li, S. Wanderling, P. Sompornpisut, E. Perozo, Structural basis of lipid-driven conformational transitions in the KvAP voltage-sensing domain. Nat Struct Mol Biol 21, 160–166 (2014).

3. B. L. Tempel, D. M. Papazian, T. L. Schwarz, Y. N. Jan, L. Y. Jan, Sequence of a probable potassium channel component encoded at Shaker locus of Drosophila. Science 237, 770–775 (1987).

4. F. Bezanilla, How membrane proteins sense voltage. Nat Rev Mol Cell Biol 9, 323–332 (2008).

5. E. Vargas et al., An emerging consensus on voltage-dependent gating from computational modeling and molecular dynamics simulations. J Gen Physiol 140, 587–594 (2012).

6. W. Treptow, M. Tarek, M. L. Klein, Initial response of the potassium channel voltage sensor to a transmembrane potential. J Am Chem Soc 131, 2107–2109 (2009).

7. M. Nishizawa, K. Nishizawa, Molecular dynamics simulation of Kv channel voltage sensor helix in a lipid membrane with applied electric field. Biophys J 95, 1729–1744 (2008).

8. V. Yarov-Yarovoy, J. Schonbrun, D. Baker, Multipass membrane protein structure prediction using Rosetta. Proteins 62, 1010–1025 (2006).

9. L. Delemotte, W. Treptow, M. L. Klein, M. Tarek, Effect of sensor domain mutations on the properties of voltage-gated ion channels: molecular dynamics studies of the potassium channel Kv1.2. Biophys J 99, L72–74 (2010).

10. Christine S. Schwaiger, P. Bjelkmar, B. Hess, E. Lindahl, 310-Helix Conformation Facilitates the Transition of a Voltage Sensor S4 Segment toward the Down State. Biophysical Journal 100, 1446–1454 (2011).

11. E. Vargas, F. Bezanilla, B. Roux, In search of a consensus model of the resting state of a voltage-sensing domain. Neuron 72, 713–720 (2011).

12. E. J. Denning, P. S. Crozier, J. N. Sachs, T. B. Woolf, From the gating charge response to pore domain movement: initial motions of Kv1.2 dynamics under physiological voltage changes. Mol Membr Biol 26, 397–421 (2009).

13. M. Nishizawa, K. Nishizawa, Coupling of S4 helix translocation and S6 gating analyzed by molecular-dynamics simulations of mutated Kv channels. Biophys J 97, 90–100 (2009).

14. C. Domene, Voltage-sensor cycle fully described. Proc Natl Acad Sci U S A 109, 8362–8363 (2012).

15. L. Delemotte, M. Tarek, M. L. Klein, C. Amaral, W. Treptow, Intermediate states of the Kv1.2 voltage sensor from atomistic molecular dynamics simulations. Proc Natl Acad Sci U S A 108, 6109–6114 (2011).

16. Y. Jiang et al., X-ray structure of a voltage-dependent K+ channel. Nature 423, 33–41 (2003).

17. J. A. Butterwick, R. MacKinnon, Solution structure and phospholipid interactions of the isolated voltage-sensor domain from KvAP. J Mol Biol 403, 591–606 (2010).

18. Z. O. Shenkarev et al., NMR structural and dynamical investigation of the isolated voltage-sensing domain of the potassium channel KvAP: implications for voltage gating. J Am Chem Soc 132, 5630–5637 (2010).

19. X. Tao, R. MacKinnon, Cryo-EM structure of the KvAP channel reveals a non-domain-swapped voltage sensor topology. Elife 8, (2019).

20. L. G. Cuello, D. M. Cortes, E. Perozo, Molecular architecture of the KvAP voltage-dependent K+ channel in a lipid bilayer. Science 306, 491–495 (2004).

21. V. Ruta, J. Chen, R. MacKinnon, Calibrated measurement of gating-charge arginine displacement in the KvAP voltage-dependent K+ channel. Cell 123, 463–475 (2005).

22. H. Biverstahl, J. Lind, A. Bodor, L. Maler, Biophysical studies of the membrane location of the voltage-gated sensors in the HsapBK and KvAP K(+) channels. Biochim Biophys Acta 1788, 1976–1986 (2009).

23. Y. Jiang, V. Ruta, J. Chen, A. Lee, R. MacKinnon, The principle of gating charge movement in a voltage-dependent K+ channel. Nature 423, 42–48 (2003).

24. C. Ahern, Stirring up controversy with a voltage sensor paddle. Trends in Neurosciences 27, 303–307 (2004).

25. C. H. Lee, R. MacKinnon, Voltage Sensor Movements during Hyperpolarization in the HCN Channel. Cell, (2019).

26. M. A. Kasimova et al., Helix breaking transition in the S4 of HCN channel is critical for hyperpolarization-dependent gating. Elife 8, (2019).

27. A. A. Gurtovenko, Asymmetry of lipid bilayers induced by monovalent salt: atomistic molecular-dynamics study. J Chem Phys 122, 244902 (2005).

28. A. A. Gurtovenko, I. Vattulainen, Calculation of the electrostatic potential of lipid bilayers from molecular dynamics simulations: methodological issues. J Chem Phys 130, 215107 (2009).

29. E. J. Denning, T. B. Woolf, Double bilayers and transmembrane gradients: a molecular dynamics study of a highly charged peptide. Biophys J 95, 3161–3173 (2008).

30. S. Chakrapani, L. G. Cuello, D. M. Cortes, E. Perozo, Structural dynamics of an isolated voltage-sensor domain in a lipid bilayer. Structure 16, 398–409 (2008).

31. Z. A. Sands, A. Grottesi, M. S. Sansom, The intrinsic flexibility of the Kv voltage sensor and its implications for channel gating. Biophys J 90, 1598–1606 (2006).

32. J. A. Freites, D. J. Tobias, Voltage Sensing in Membranes: From Macroscopic Currents to Molecular Motions. J Membr Biol 248, 419–430 (2015).

33. J. A. Freites, E. V. Schow, S. H. White, D. J. Tobias, Microscopic origin of gating current fluctuations in a potassium channel voltage sensor. Biophys J 102, L44–46 (2012).

34. S. F. Altschul et al., Gapped BLAST and PSI-BLAST: a new generation of protein database search programs. Nucleic Acids Res 25, 3389–3402 (1997).

35. S. I. B. S. I. o. B. Members, The SIB Swiss Institute of Bioinformatics’ resources: focus on curated databases. Nucleic Acids Res 44, D27–37 (2016).

36. H. M. Berman et al., The Protein Data Bank. Nucleic Acids Res 28, 235–242 (2000).

37. S. Jo, J. B. Lim, J. B. Klauda, W. Im, CHARMM-GUI Membrane Builder for mixed bilayers and its application to yeast membranes. Biophys J 97, 50–58 (2009).

38. W. L. Jorgensen, J. Chandrasekhar, J. D. Madura, R. W. Impey, M. L. Klein, Comparison of simple potential functions for simulating liquid water. The Journal of Chemical Physics 79, 926–935 (1983).

39. A. P. Demchenko, S. O. Yesylevskyy, Nanoscopic description of biomembrane electrostatics: results of molecular dynamics simulations and fluorescence probing. Chem Phys Lipids 160, 63–84 (2009).

40. C. Kutzner, H. Grubmuller, B. L. de Groot, U. Zachariae, Computational electrophysiology: the molecular dynamics of ion channel permeation and selectivity in atomistic detail. Biophys J 101, 809–817 (2011).

41. M. Casciola, D. Bonhenry, M. Liberti, F. Apollonio, M. Tarek, A molecular dynamic study of cholesterol rich lipid membranes: comparison of electroporation protocols. Bioelectrochemistry 100, 11–17 (2014).

42. A. Polak et al., On the electroporation thresholds of lipid bilayers: molecular dynamics simulation investigations. J Membr Biol 246, 843–850 (2013).

43. D. Van Der Spoel et al., GROMACS: fast, flexible, and free. J Comput Chem 26, 1701–1718 (2005).

44. A. D. MacKerell et al., All-atom empirical potential for molecular modeling and dynamics studies of proteins. J Phys Chem B 102, 3586–3616 (1998).

45. A. D. Mackerell, Jr., M. Feig, C. L. Brooks, 3rd, Extending the treatment of backbone energetics in protein force fields: limitations of gas-phase quantum mechanics in reproducing protein conformational distributions in molecular dynamics simulations. J Comput Chem 25, 1400–1415 (2004).

46. J. B. Klauda et al., Update of the CHARMM all-atom additive force field for lipids: validation on six lipid types. J Phys Chem B 114, 7830–7843 (2010).

47. H. J. C. Berendsen, J. P. M. Postma, W. F. Vangunsteren, A. Dinola, J. R. Haak, Molecular-Dynamics with Coupling to an External Bath. Journal of Chemical Physics 81, 3684–3690 (1984).

48. G. Bussi, D. Donadio, M. Parrinello, Canonical sampling through velocity rescaling. J Chem Phys 126, 014101 (2007).

49. B. Hess, H. Bekker, H. J. C. Berendsen, J. G. E. M. Fraaije, LINCS: A linear constraint solver for molecular simulations. Journal of Computational Chemistry 18, 1463–1472 (1997).

50. U. Essmann et al., A smooth particle mesh Ewald method. The Journal of Chemical Physics 103, 8577–8593 (1995).

51. S. Jo, T. Kim, W. Im, Automated builder and database of protein/membrane complexes for molecular dynamics simulations. PLoS One 2, e880 (2007).

52. F. E. Regnier, W. Cho, in Proteomic and Metabolomic Approaches to Biomarker Discovery. (2013), pp. 197–224.

53. K. Vanommeslaeghe et al., CHARMM general force field: A force field for drug-like molecules compatible with the CHARMM all-atom additive biological force fields. J Comput Chem 31, 671–690 (2010).

54. J. J. Irwin, T. Sterling, M. M. Mysinger, E. S. Bolstad, R. G. Coleman, ZINC: a free tool to discover chemistry for biology. J Chem Inf Model 52, 1757–1768 (2012).

55. O. Livnah, E. A. Bayer, M. Wilchek, J. L. Sussman, Three-dimensional structures of avidin and the avidin-biotin complex. Proc Natl Acad Sci U S A 90, 5076–5080 (1993).

